# A GPI-anchored Ly6/uPAR superfamily gene *belly roll* is expressed in multiple peptidergic neurons in *Drosophila melanogaster* larvae

**DOI:** 10.64898/2026.02.27.708413

**Authors:** Yuma Tsukasa, Tadashi Uemura, Tadao Usui

**Affiliations:** Graduate School of Biostudies, Kyoto University; Center for Living Systems Information Science (CeLiSIS), Kyoto University

**Keywords:** Ly6/uPAR superfamily, Belly roll, Drosophila, Peptidergic neurons, Single-cell RNA sequencing, T2A-GAL4 system

## Abstract

The Lymphocyte antigen-6 (Ly6)/urokinase-type plasminogen activator receptor (uPAR) superfamily (LU super family) of proteins are involved in diverse biological processes. In *Drosophila melanogaster*, members of the LU superfamily have undergone lineage-specific gene duplication and acquired specialized functions in distinct tissues. A glycosylphosphatidylinositol (GPI)-anchored LU family protein Belly roll (Bero) has recently been shown to regulate larval escape behavior; however, its cellular expression profile and potential roles remain incompletely understood. In this study, we generated a *bero-GAL4*^*T2A*^ transgenic line to delineate endogenous *bero* expression. This analysis revealed that *bero* is expressed in the peptidergic neurons in the central nervous system (CNS) that had not been documented in previous studies, as well as in the peripheral nervous system (PNS) and non-neuronal tissues, such as the anal pad and epidermis. Reanalysis of publicly available single-cell RNA sequencing (scRNA-seq) datasets demonstrated that *bero* is expressed in several peptidergic neurons. These findings suggest that Bero is specifically expressed in diverse peptidergic neurons and may play important roles in coordinating hormonal and neural regulation in *D. melanogaster*.

## 1 Introduction

The Lymphocyte antigen-6 (Ly6)/urokinase-type plasminogen activator receptor (uPAR) superfamily of proteins are conserved from vertebrates to insects (Leth et al., 2019). Most of LU proteins are GPI-anchored to the outer leaflet of cell membrane and involved in diverse processes such as immune regulation, cell-cell adhesion, sperm-egg recognition, cell migration and neuromodulation (Loughner et al., 2016; Miwa et al., 2012). In *Drosophila melanogaster*, the LU gene family is predicted to have expanded through multiple gene duplication and acquired diverse physiological functions in various tissues (Baudouin-Gonzalez et al., 2017; Hijazi et al., 2009; Moussian et al., 2005; Tanaka et al., 2015). For example, *boudin* and *coiled* are expressed in the embryonic tracheal epithelium and is required for septate junctions’ integrity (Hijazi et al., 2011, 2009; Nilton et al., 2010). Furthermore, *sleepless* is expressed broadly in the adult brain and regulates sleep by binding to either an A-type potassium ion channel Shaker or nicotinic acetylcholine receptors (Koh et al., 2008; Wu et al., 2016, 2014; Wu et al., 2010).

Our recent study has revealed that *belly roll* (*bero*)/*CG9336* is engaged in neuromodulation of the escape behavior in *Drosophila* larvae (Li et al., 2023). *bero* is one of the most recently duplicated LU family genes, expressed in diverse organs including the central nervous system (CNS), intestine, and heart during embryonic development (Baudouin-Gonzalez et al., 2017; Tanaka et al., 2015). In larvae, *bero* is expressed in several groups of neurons and glia in the CNS such as abdominal leucokinin-producing neurons (ABLK neurons), insulin-like peptides-producing cells (IPC), a subset of eclosion hormone-producing neurons (EH neurons), Capability peptides-producing neurons (CAPA neurons) and midline glia (MG) (Table 1; Baudouin-Gonzalez et al., 2017; Li et al., 2023). In ABLK neurons, Bero is suggested to regulate neurohormone secretion through modulation of neural activity (Li et al., 2023). It is noteworthy that large population of *bero*-expressing cells coexpresses *dimmed* that encodes a bHLH transcription factor enriched in peptidergic neurons (Li et al., 2023). Based on these observations, it was hypothesized that Bero is widely implicated in the secretion of neuropeptides and neurohormones. However, many *bero*^+^ peptidergic cells remain unidentified.

**Table 1.** *bero*^*+*^ Cells in larvae.

| Cell type | Driver | Source | References |
| --- | --- | --- | --- |
| IPC | ilp2-GAL4 | BDSC: 37516 | Li et al., 2023 |
| EH neurons | Eh.2.4-GAL4 | BDSC: 6301 | Li et al., 2023 |
| CAPA neurons |  |  | Baudouin-Gonzalez et al., 2017 |
| ABLK neurons | Lk-GAL4 | BDSC: 51993 | Li et al., 2023 |
| GPA2/GPB5 neurons | Gpb5-GAL4 | Sellami et al., 2011 | This study |
| OK neurons | Ok-GAL4 | Chen et al., 2015 | This study |
| Proc+ motor neurons | Proc-GAL4 | BDSC: 51972 | This study |
| Midline Glia |  |  | Baudouin-Gonzalez et al., 2017 |
| Eye disc glia |  |  | Baudouin-Gonzalez et al., 2017 |
| Garland cells |  |  | Baudouin-Gonzalez et al., 2017 |
| Hindgut |  |  | Baudouin-Gonzalez et al., 2017 |
| Heart |  |  | Baudouin-Gonzalez et al., 2017 |
| Bolwig's organ |  |  | Baudouin-Gonzalez et al., 2017 |

*Note:* BDSC: Bloomington Drosophila stock center

In this study, we generated a *bero-GAL4*^*T2A*^ transgenic line to help detect endogenous *bero* expression more extensively. This line revealed that *bero* is expressed in the several peptidergic neurons in central nervous system (CNS) that had not been documented in previous studies, as well as in peripheral nervous system (PNS) and non-neuronal tissues, such as the anal pad and epidermis. A reanalysis of publicly available single-cell RNA sequencing (scRNA-seq) datasets and histological analyses using confocal laser scanning microscopy have revealed that several peptidergic neurons express *bero* gene (Table 1). In addition, we found that such *bero+* peptidergic neurons coexpress multiple neuropeptides or neurohormones (Table 2). These results suggest that Bero may play an important role in physiological processes of several peptidergic neurons as well as in neuromodulation of ABLK neurons.

**Table 2.**

| Cell type | Transmitters | References |
| --- | --- | --- |
| IPC | Dilp1,2,3,5, Dsk | see the comprehensive review Nässel et al., 2020 |
| EH neurons | EH | see the comprehensive review Nässel et al., 2020 |
| CAPA neurons | CAPA | see the comprehensive review Nässel et al., 2020 |
| ABLK neurons | Lk, Dh44, TA and/or OA, Dh31* | Zandawala et al. (2018); Li et al. (2023); This study |
| GPA2/GPB5 neurons | Gpa2, Gpb5, Dh31*, AstA* | Sellami et al. (2011); This study |
| OK | Orcokinin A | Chen et al. (2015) |

*Note:* \* Detected by the *GAL4<sup>T2A</sup>* systems or scRNA-seq analysis

## 2 Result

### 2.1 Expression pattern of *bero-GAL4*^*T2A*^ driver line

To thoroughly investigate the endogenous expression of the *bero* gene, we employed the T2A-GAL4 reporter system (Figure 1A). In this system, following the initial translation of *bero* coding sequence, a ribosomal skipping is induced by the viral T2A sequence, resulting in the subsequent translation of GAL4 as an independent polypeptide from the same messenger RNA. The endogenous expression pattern of *bero* was analyzed by crossing *bero-GAL4*^*T2A*^ line with *UAS-reporter* lines (Figure 1A). As previously reported (Li et al., 2023), *bero*-expressing cells were observed in several subpopulations in peptidergic neurons in CNS (Figure 1B-a). Imaging from the lateral body wall revealed that some epidermal cells and peripheral nerves were also labeled by *bero-GAL4*^*T2A*^ (Figure 1B-b). In addition, the anal pads were strongly labeled (Figure 1B-c).

**Figure 1.**
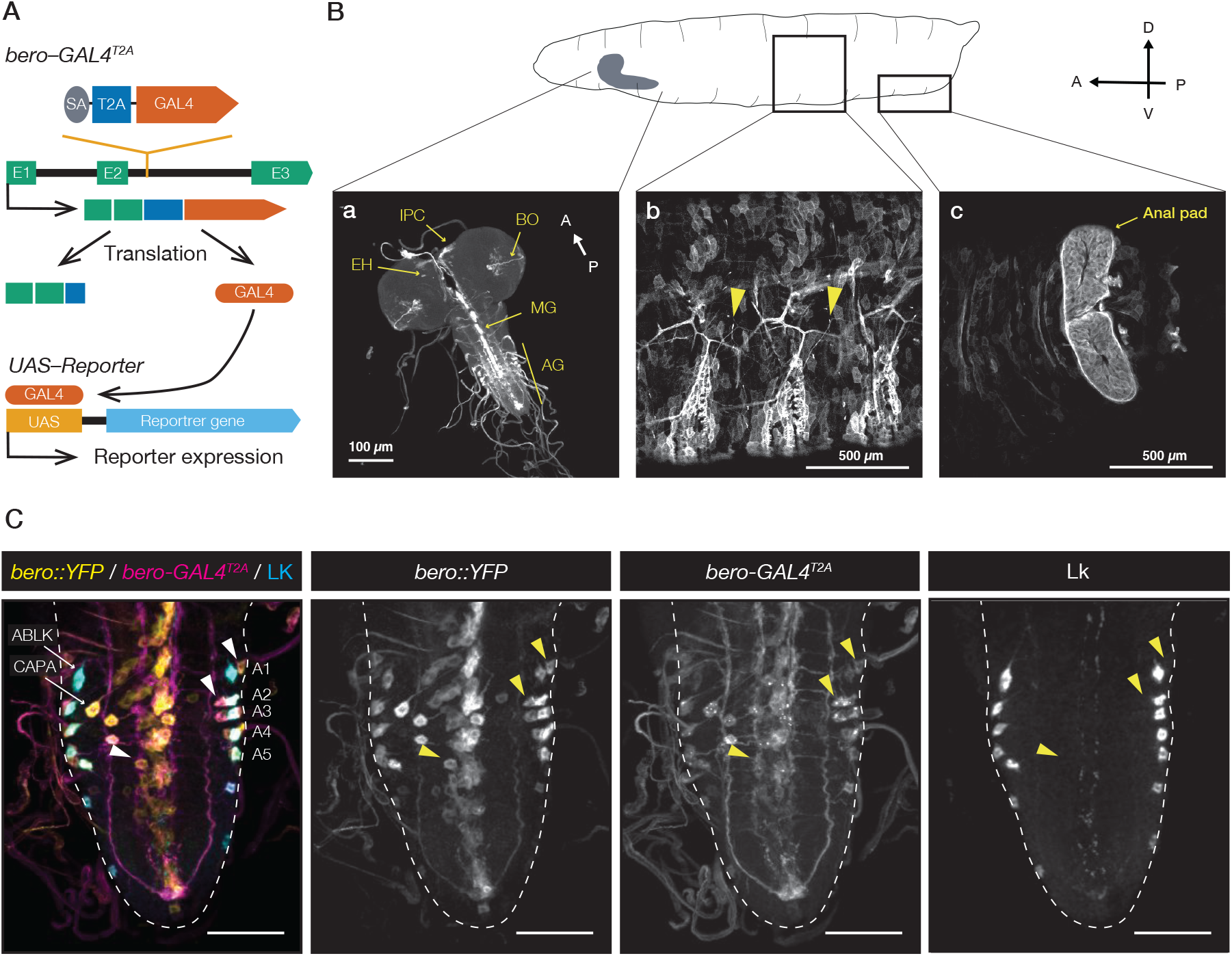
Endogenous expression patterns of *bero* in *Drosophila* larvae. (A) Overview of a method for visualizing *bero* expression patterns using *bero-GAL4*^*T2A*^. (B) Confocal microscopy images of the expression pattern of *bero-GAL4*^*T2A*^ in CNS (a), lateral body wall (b) and anal pad (c) of *Drosophila* larvae. IPC: Insulin-like peptide producing cells; EH: Eclosion hormone producing neurons; BO: Bolwig organs; MG: Midline glia; AG: abdominal ganglion. (C) Confocal microscope stack images of *bero-GAL4*^*T2A*^ (magenta) or *bero::YFP* labeled cells (yellow) and ABLK neurons (cyan) in the CNS. Arrows indicate previously identified *bero*^+^ cells, while triangles indicate unknown *bero*^+^ cells. Scale bars: 50 μm.

A previous study has demonstrated that ABLK neurons are labeled by a protein-trap strain *bero::YFP* (Lowe et al., 2014; Li et al., 2023). We asked whether *bero-GAL4*^*T2A*^ and *bero::YFP* lines express the transgenes in the same fashion, and then aimed to compare their expression patterns in ABLK neurons (Figure 1C). Consistent with previous studies (Baudouin-Gonzalez et al., 2017; Li et al., 2023), *bero::YFP* labeled ABLK neurons and CAPA neurons, while *bero-GAL4*^*T2A*^ express *GAL4* in the same neurons (Figure 1C). We then noticed previously unidentified *bero*^+^ cells with large cell body along the midline glia of the abdominal ganglion (Figure 1C). Furthermore, several unidentified *bero*^+^ cells were also detected in proximity to the ABLK neurons in some abdominal neuromeres (A1–A5; Figure 1C). These two independent cell-labeling tools allowed us to robustly highlight *bero*-expressing cells without misidentifying spurious gene expression. These results indicated that the *bero-GAL4*^*T2A*^ line established in this study is a valuable tool for the analysis of *bero* expression patterns.

### 2.2 scRNA-seq analysis revealed the expression of *bero* in midline glia, subpopulations of peptidergic neurons

To identify *bero*^+^ cells, we reanalyzed publicly available single-cell RNA sequencing (scRNA-seq) datasets (Nguyen et al., 2024). These datasets were obtained from dissociated cells of the ventral nerve cord (VNC) in *Drosophila* third instar larvae and classified into 18 cell-types (Figure 2A; Nguyen et al., 2024). The quantification of *bero* expression levels in each cell and the subsequent plotting by cluster revealed that *bero* is highly expressed in midline glia and in a subset of motor neurons/neurosecretory cells (MG in Fig. 2B and C). These results are consistent with previous reports

**Figure 2.**
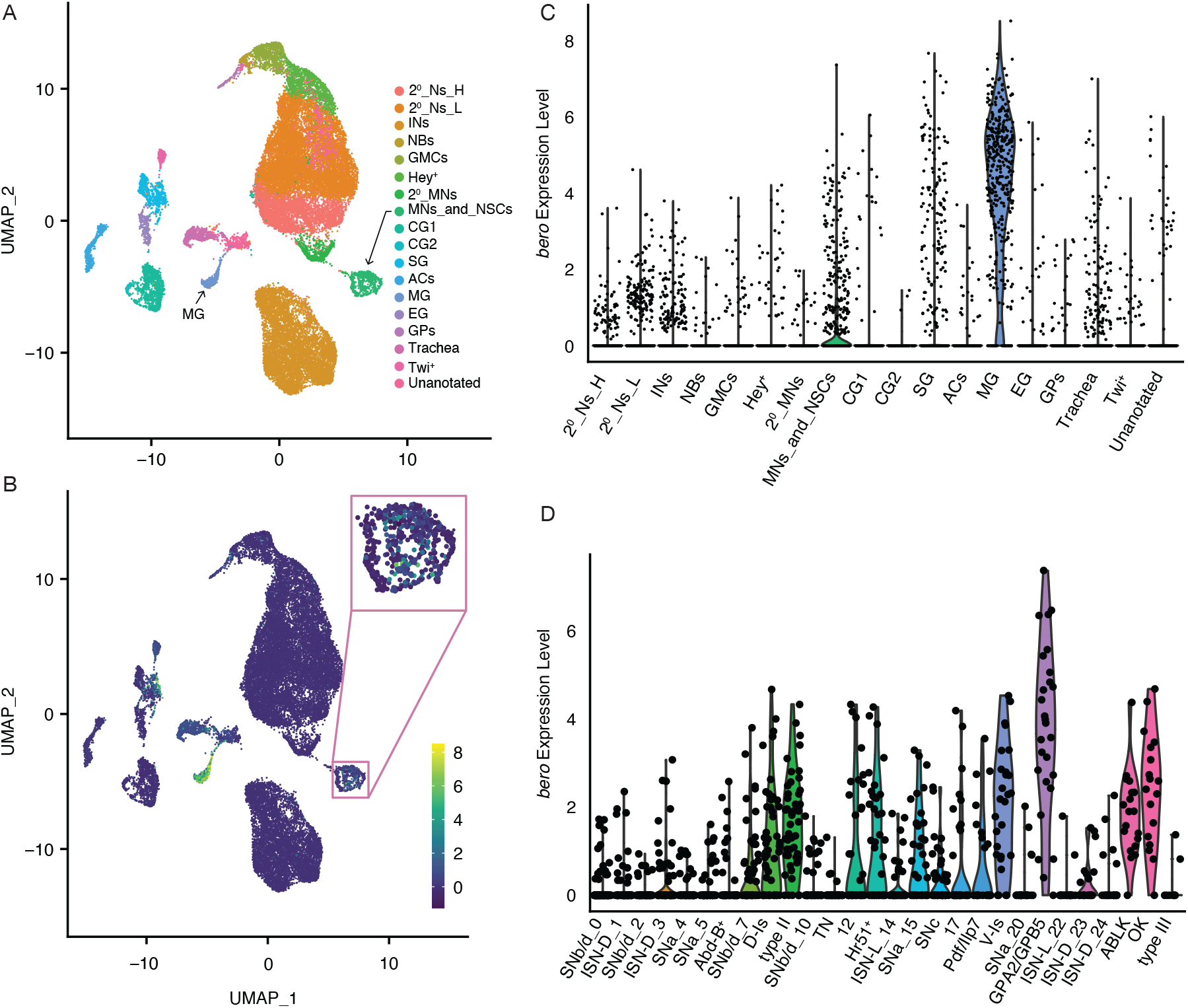
Identification of *bero*^+^ subpopulations via scRNA-seq analysis. (A) Two-dimensional representation of all cells (UMAP). The cluster classification was reported in Nguyen et al., 2024. 2^0^_Ns_H: Secondary neurons (high lmp); 2^0^_Ns_L: Secondary neurons (low lmp); Ins: Inter neurons; NBs: Neuroblasts; GMCs: Ganglion mother cells; Hey^+^: Hey^+^ newborn neurons; 2^0^_MNs: Secondary motor neurons; CG1: Cortex glia 1; CG2: Cortex glia 2; SG: Subperineurial glia; ACs: Astrocytes; MG: Midline glia; EG: Ensheathing glia; GPs: Glia precursors. (B-C) Density plots (B) and violin plots (C) showing that *bero* transcripts were highly expressed in midline glia (MG) and motor neurons and neurosecretory cells (MNs_and_NSCs). (D) Violin plot showing the expression levels of bero in a subset of *twit* (MNs_and_NSCs marker) positive cells.

(Baudouin-Gonzalez et al., 2017; Li et al., 2023) and the analysis of *bero-GAL4*^*T2A*^ expression pattern (Figure 1Ba and C).

To further investigate *bero*^+^ cell types, we examined which subpopulations within each motor neuron/neurosecretory cell cluster express *bero* (Figure 2D). This analysis revealed that *bero* was expressed in Dorsal-Is (D-Is) and Ventral-Is (V-Is) motor neurons that produce the neuropeptide Proc, Type II motor neurons, Gpa2/ Gpb5-producing neurons (GPA2/GPB5 neurons), ABLK neurons, and Ok-producing neurons (OK neurons). These results support the previous prediction that several populations of *bero*^*+*^ cells are peptidergic neurons (Li et al., 2023). The *proc*^+^ D-Is and V-Is motor neurons, whose cell bodies are located near the midline (Taylor et al., 2004), might represent previously unidentified *bero*^+^ cells that were labeled by *bero-GAL4*^*T2A*^. Furthermore, given that GPA2/GPB5 neurons and OK neurons are located in A1-A4 and A1-A5 segments, respectively, and differentiate from a cell lineage shared with ABLK (Díaz-de-la-Peña et al., 2020; Sellami et al., 2011), these peptidergic neurons were also strong candidates for the uncharacterized *bero*^+^ cells.

### 2.3 *bero* was expressed in *proc*^+^ motor neurons, GPA2/GPB5 neurons and OK neurons

To validate the *bero*^+^ cells candidates estimated by scRNA-seq analysis, we examined the overlaps on a cell-by-cell basis between the expression pattern of cell type-specific GAL4 drivers and that of *bero::YFP* protein-trap reporter. Among the cells that were labeled with *proc-GAL4*, several subpopulations were also labeled with *bero::YFP* (Fig. 3A-a). The subpopulations whose cell bodies are located near the midline were considered to be *proc*^+^ motor neurons (Fig. 3A-b; Nguyen et al., 2024). In addition, some *proc*^+^∩*bero*^+^ cells of unknown identity were also observed (Fig.3A-c and 3A-d). We reanalyzed the scRNA-seq dataset of *proc*^+^ sorted VNC cells and found that *bero* was expressed not only in D-Is and V-Is but also in *Syncrip*^+^ (*Syp*^+^) motor neurons and unannotated cells (Fig. 3B). These cell populations identified by scRNA-seq analysis may be lateral *proc*^+^∩ *bero*^+^ neurons. A subsequent comparison of the expression patterns of *Gpb5-GAL4* or *Ok-GAL4* and *bero::YFP* revealed overlaps between them (*Gpb5*^+^∩*bero*^+^ or *Ok* ^+^∩*bero*^+^ cells, respectively; Fig. 3C and D). Taken together, these findings indicate that *bero* is expressed in several peptidergic neurons, including *proc*^+^ motor neurons, GPA2/GPB5 neurons, and OK neurons.

**Figure 3.**
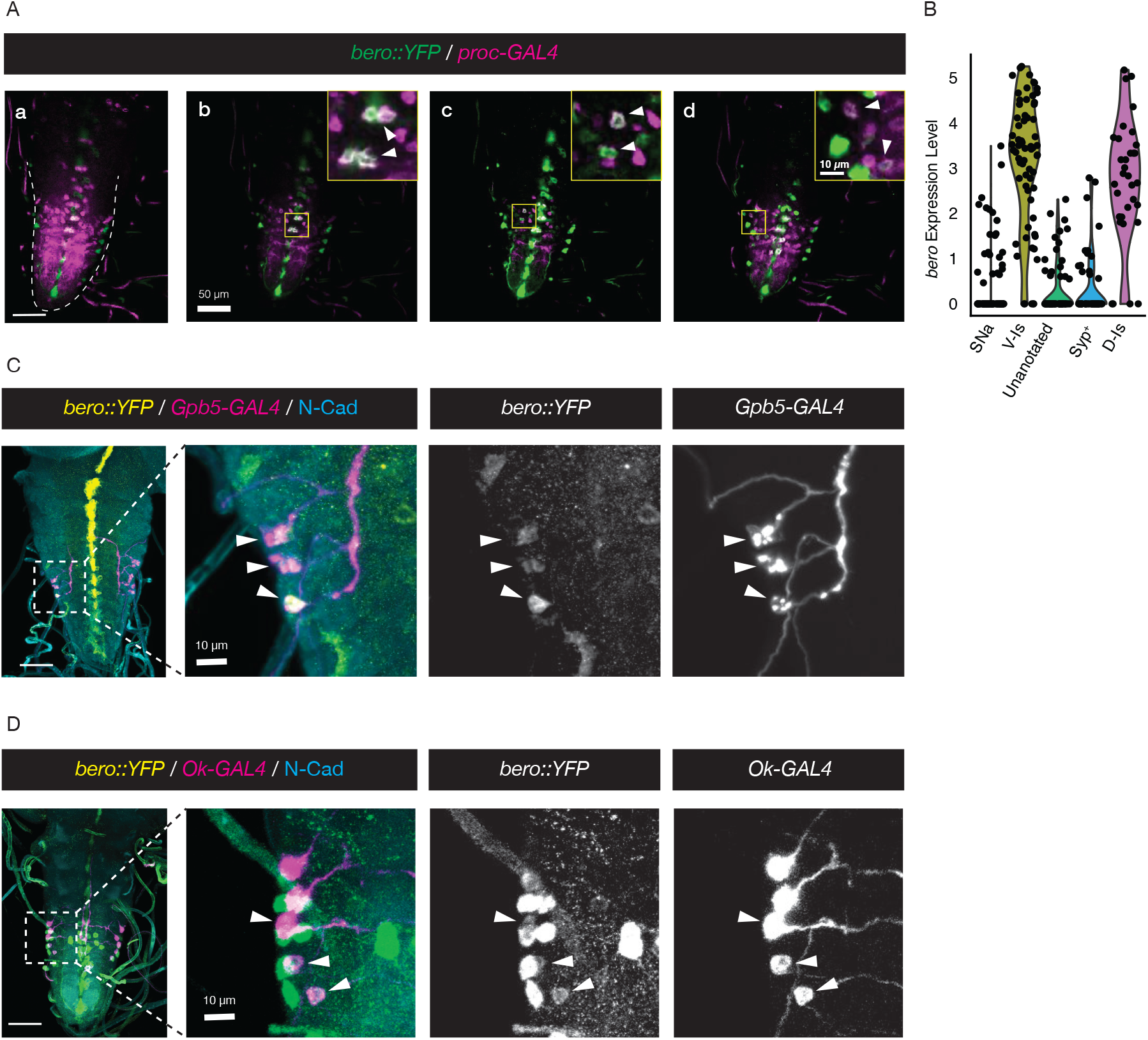
*bero* is expressed in several subpopulations of neurosecretory cells. (A) Confocal microscope stacks (a) or single sections (b-d) showing *bero::YFP* labeled cells (green) and *Proc-GAL4* labeled cells (magenta) in the CNS. Scale bars: 50 μm. (B) A violin plot of *bero* transcripts in *proc*^+^ sorted VNC cells. (C) Confocal microscope stacks showing Bero::YFP labeled cells (yellow) and *Gpb5-GAL4* labeled cells (magenta) in the CNS. Scale bars: 50 μm or 10 µm. (D) Confocal microscope stacks showing *bero::YFP* labeled cells (yellow) and *Ok-GAL4* labeled cells (magenta) in the CNS. Scale bars: 50 μm or 10 µm.

### 2.4 Coexpression of neuropeptides in *bero*^+^ peptidergic neurons

It is well established that most peptidergic neurons co-secrete multiple neuropeptide and/or neurohormones (Jiménez et al., 2006; Kahsai et al., 2010; Nässel & Zandawala, 2019; S. R. Taylor et al., 2021). For instance, ABLK neurons coproduce the Diuretic hormone 44 (Dh44) in conjunction with Lk (Nässel, 2021). To explore the coexpression of neuropeptides and/or neurohormones in *bero*^+^ peptidergic neurons, we conducted another reanalysis of the scRNA-seq datasets. Consistent with our aforementioned result, D-Is and V-Is motor neurons coexpressed *proc* (Fig.4A). Furthermore, in consistent with a previous study (Nguyen et al., 2024), it has been suggested that GPA2/GPB5 neurons coexpressed *Allatostatin A* (*AstA*). In addition, we discovered that GPA2/GPB5 neurons and ABLK neurons coexpressed *Diuretic hormone 31* (*Dh31*).

To further validate the expression of neuropeptides and/or neurohormones, the following four T2A-GAL4 lines were utilized to label specific peptidergic cell types: *AstA-GAL4*^*T2A*^, *Dh44*-*GAL4*^*T2A*^, *Dh31*^*AC*^*-GAL4*^*T2A*^ and *Dh31*^*D*^*-GAL4*^*T2A*^. We then found that *AstA-GAL4*^*T2A*^ labeled *bero*^+^∩*Lk*^−^ cells in the close vicinity of ABLK neurons (Fig. 4B). Consistent with prior reports employing a distinct *Dh44-GAL4* driver line (*Dh44-GAL4*^*R65C11*^; Zandawala, Marley, et al., 2018), the *Dh44-GAL4*^*T2A*^ line labeled all ABLK neurons from A1 to A7 (Fig. 4C). The two distinct *Dh31-GAL4*^*T2A*^ lines were generated based on different designs (Deng et al., 2019): one was engineered to express *GAL4* as a separate polypeptide downstream of the coding sequence of RA and RC isoforms of *Dh31* (*Dh31*^*AC*^*-GAL4*^*T2A*^), while the other was to express *GAL4* downstream of RD isoform (*Dh31*^*D*^*-GAL4*^*T2A*^). Both *Dh31-GAL4*^*T2A*^ were found to label a specific subset of ABLK neurons (Fig. 4D and E); *Dh31*^*AC*^*-GAL4*^*T2A*^ was predominantly expressed in A6 and A7 neuromeres, whereas *Dh31*^*D*^*-GAL4*^*T2A*^ was only in A7 (Fig.4F). In summary, the peptidergic neurons that express *bero* also produce and secrete a variety of neuropeptides and/or neurohormones (Table 2).

**Figure 4.**
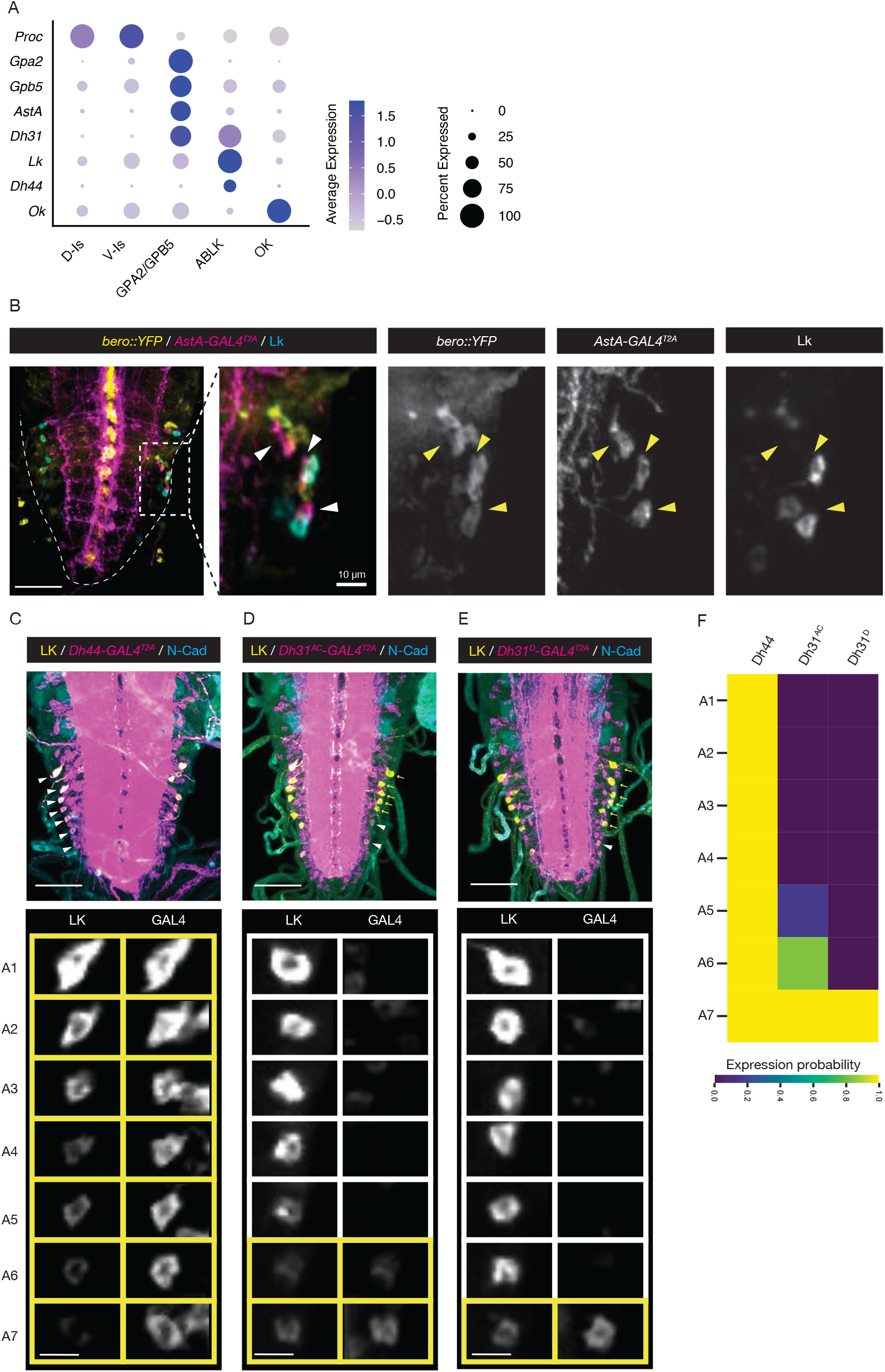
Expression profiles of neuropeptide and/or neurohormones in *bero*^+^ neurosecretory cells. (A) A dot plot of neuropeptides and neurohormones in D-Is motor neurons, V-Is motorneurons, GPA2/GPB5 neurons, ABLK neurons and OK neurons. (B) Confocal microscope stacks showing *bero::YFP* labeled cells (yellow), *AstA-GAL4*^*T2A*^ labeled cells (magenta) and ABLK neurons (cyan) in the CNS. Scale bars: 50 μm or 10 µm. (C-E) Confocal microscope stacks showing ABLK neurons (yellow) and *Dh44-GAL4*^*T2A*^ labeled cells (C; magenta), *Dh31*^*AC*^*-GAL4*^*T2A*^ labeled cells (D; magenta) or *Dh31*^*D*^*-GAL4*^*T2A*^ labeled cells (E; magenta). Scale bars: 50 μm or 10 µm. (F) A heat map showing expression probability of *Dh44, Dh31*^*AC*^ and *Dh31*^*D*^ in ABLK neurons.

Our scRNA-seq reanalyses suggested that GPA2/GPB5 neurons express *AstA* and *Dh31*, but these finding have not been directly confirmed by independent approaches using GAL4^T2A^ systems. However, *AstA-GAL4*^*T2A*^-labeled *bero*^+^∩*Lk*^−^ cells have been localized in close proximity to ABLK neurons (Figure 4B), and this anatomical feature was consistent with the characteristics of GPA2/GPB5 neurons (Sellami et al., 2011). Similarly, GPA2/GPB5 neurons may have been included among the *Dh31*^+^ cells in proximity to ABLK neurons.

## 3 Discussion

In this study, we took a comprehensive strategy and revealed that the larval unidentified neurons expressing *bero* are the peptidergic neurons including *proc*^+^ motor neurons, GPA2/GPB5 neurons, and OK neurons (Fig. 5). Furthermore, we examined coexpressing neuropeptides and/or neurohormones in *bero*^+^ peptidergic neurons. Although it has been established that ABLK neurons coexpress *Dh44* in conjunction with *Lk*, we have further revealed that a specific subset of ABLK neurons coexpress *Dh31* (Fig. 5). Moreover, scRNA-seq analysis indicated that GPA2/GPB5 neurons potentially express *AstA* and *Dh31* (Fig. 5).

**Figure 5.**
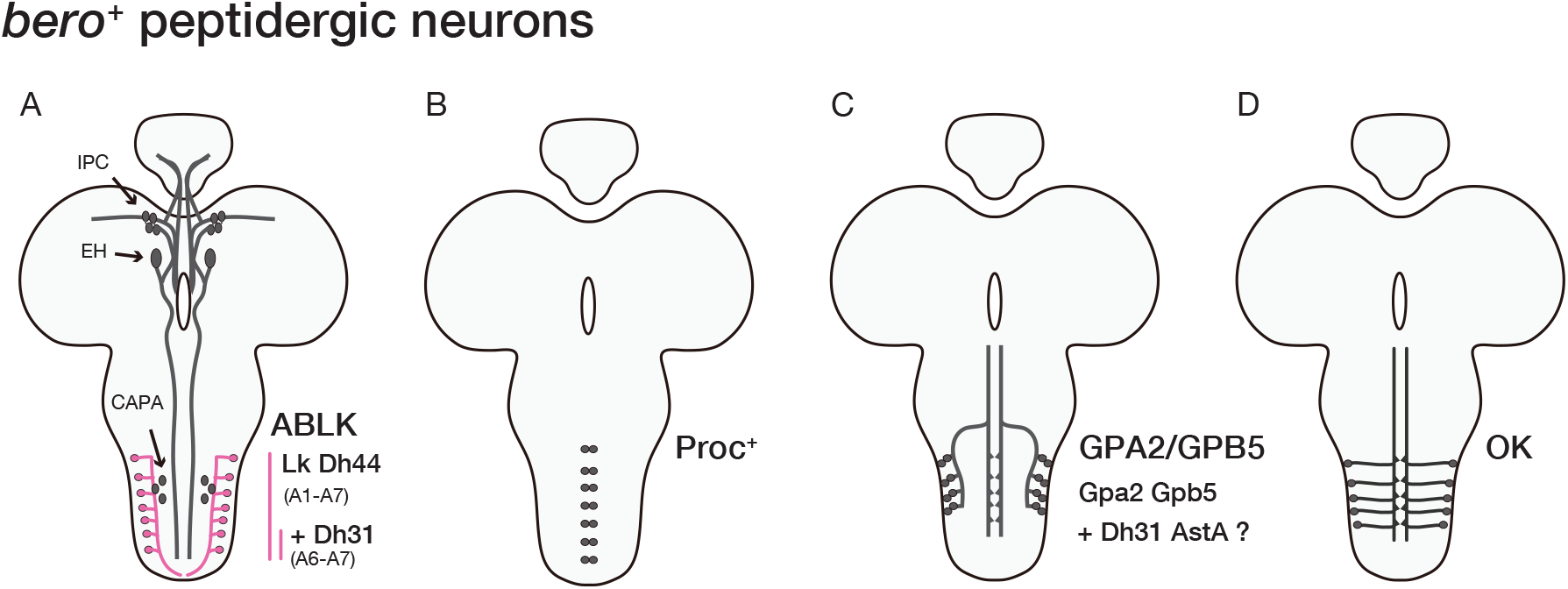
Schematic of the *bero*-expressing peptidergic neurons. (A) The *bero*^+^ peptidergic neurons that were identified in previous research (Baudouin-Gonzalez et al., 2017; Li et al., 2023). ABLK neurons coexpressed Dh31. (B-D) The *bero*^+^ peptidergic neurons that were identified in this study. GPA2/GPB5 neurons possibly expressed Dh31 and AstA.

The small size of the central nervous system in *Drosophila* larvae, comprising approximately 10,000 cells, enables highly specific annotation of cell type-specific signaling molecules. While connectome information for *Drosophila* larvae has been published in recent years (Hückesfeld et al., 2021; Winding et al., 2023), establishing a neuropeptidergic connectome which represents synaptic transmission across different layers within neural circuitry must be essential for a deeper understanding of brain function, as has already been achieved for nematodes (Ripoll-Sánchez et al., 2023). The extensive combination of scRNA-seq analyses and a specific labeling system using *T2A-GAL4* employed in this study may facilitate the establishment of a neuropeptidergic connectome in *Drosophila* larvae (Deng et al., 2019; Kondo et al., 2020; Mao et al., 2024).

In ABLK neurons, Bero has been shown to inhibit neurosecretion by regulating neural activity (Li et al., 2023). Given the expression of Bero in a variety of peptidergic neurons, it is possible that the inhibition by Bero may represent a universal mechanism in cells other than ABLK neurons. Notably, the some neuropeptide and/or neurohormones produced in *bero*^+^ cells modulate organismal nutritional status (Al-Anzi et al., 2010; Ohashi & Sakai, 2018; Zandawala, Yurgel, et al., 2018) and/or water and ion homeostasis (e.g., CAPA, Lk, Gpa2/Gpb5, Dh31; Cabrero et al., 2002; Chu et al., 2024; Liu et al., 2015; Okamoto et al., 2026; Paluzzi et al., 2014; Yoon et al., 2023). The physiological relevance of Bero in the maintenance of internal homeostasis remains to be elucidated; however, based on the known function of the *bero*^+^ cells, it is likely that Bero plays important roles in homeostasis.

## 4 Experimental Procedures

### 4.1 Fly husbandry

*D. melanogaster* strains were reared at 25°C and 75-80% humidity with a 12 hr light/dark cycle on standard fly food as previously described (Watanabe et al., 2019). The fly strains used were as follows: *bero-GAL4*^*T2A*^ (this study); *bero::YFP* (Kyoto Drosophila stock center: 115180), *Lk-GAL4* (BDSC: 51993), *Gpb5-GAL4* (Sellami et al., 2011); *Ok-GAL4* (Chen et al., 2015; Gift from Jae Young Kwon); *Proc-GAL4* (BDSC: 51972); *AstA-GAL4*^*T2A*^ (BDSC: 84593); *Dh44-GAL4*^*T2A*^ (BDSC: 84627); *Dh31*^*AC*^*-GAL4*^*T2A*^ (BDSC: 84623); *Dh31*^*D*^*-GAL4*^*T2A*^ (BDSC: 84624); *UAS-myr::GFP* (BDSC: 32197); *UAS-hCD4::tdTomato* (BDSC: 35837); *UAS-myr::tdTomato* (BDSC: 32221).

### 4.2 Generation of *bero-GAL4*^***T2A***^

*bero-GAL4*^*T2A*^ strain was generated by inserting pBS-KS-attB2-SA(1)-T2A-Gal4-Hsp70 (Addgene: #62897) into MiMIC landing site of *bero*^MI03849^ strain (BDSC: 36397; WellGenetics Inc., Taipei, Taiwan) by φC31-mediated genomic integration, as previously described (Diao et al., 2015).

### 4.3 Immunohistochemistry and confocal imaging

For the detection of LK peptide or *D*N-Cadherin in the larval CNS, wandering third instar larvae were dissected in PBS (∼303±1.53 mOsm/kg, which is very close to the osmolality of *Drosophila* larvae hemolymph; Kurio et al., 2024) and fixed with 4% formaldehyde in PBS at 4°C overnight. After fixation, the dissected CNS were washed five times with PBST (PBS containing 0.3% Triton X-100) and blocked in PBST containing 2% BSA (filtrated with 0.22 μm filter) for 30 min at room temperature. The samples were then incubated with rabbit polyclonal anti-LK (1:500; Ohashi and Sakai, 2018) or rat monoclonal anti-N-cadherin (1:100; DN-Ex #8, Developmental Studies Hybridoma Bank [DSHB], Iowa City, IA) at 4°C for 3 days. After five washes with PBST, the samples were incubated with the Alexa Fluor 405-conjugated goat polyclonal anti-rabbit IgG (H+L) (1:500; A-31556, Thermo Fisher Scientific) for 2 or 3 hours at room temperature. After further washes, the samples were mounted with ProLong Glass Antifade Mountant (Thermo Fisher, Carlsbad, CA, USA). Images were taken with a Nikon C1Si confocal microscope and processed with Fiji (ImageJ, NIH, Bethesda, MD, USA). The data processing and visualization of the heatmap (Fig. 4F) were performed using the pandas (ver.1.1.2) and seaborn (ver.0.12.2) in Python (ver.3.7.7).

### 4.4 scRNA-seq data analysis

Previously published third instar larval CNS scRNA-Seq data (GSE235231; Nguyen et al., 2024) was employed. The data processing and visualization were performed using the Seurat package (ver.3.2.2; Stuart et al., 2019) and ggplot2 (ver.3.3.2) in R (ver.3.6.1).

## Conflict of Interest

The authors declare no competing or financial interests.

## Funding

The Japan Society for the Promotion of Science 21K06264 (TUsui)

The Kyoto University Foundation (TUsui)

ISHIZUE 2024 of Kyoto University (TUsui)

The Japan Society for the Promotion of Science 22KJ1999 (YT)

The Uehara Memorial Foundation (TUsui)

## Author contributions

Conceptualization, Y.T., T.Us.; Methodology, Y.T., T.Us.; Software, Y.T.; Validation, Y.T., T.Ue., T.Us.; Formal analysis, Y.T.; Investigation, Y.T., T.Us; Resources, T.Us.; Data curation, Y.T.; Writing - original draft, Y.T., T.Us.; Writing - review & editing, Y.T., T.Ue., T.Us.; Visualization, Y.T., T.Us.; Supervision, T.Us.; Project administration, T.Us.; Funding acquisition, Y.T., T.Us.

## Data availability

All relevant data can be found within the article and its supplementary information.

## Acknowledgments

We would like to thank M. Futamata, S. Oki, H. Imai, R. Muraki, and Y. Niitani for excellent technical assistance; members of the Uemura lab for experimental advice and discussions; Takaomi Sakai (Tokyo Metropolitan Univ.) for the anti-LK primary antibody; Pilar Herrero for *Gpb5-GAL4*; Jae Young Kwon for *Ok-GAL4*; Bloomington *Drosophila* Stock Center, Kyoto Drosophila Stock Center and Korea Drosophila Resource Center for providing fly stocks. We also thank FlyBase and the *Drosophila* Genomics Resource Center.

## Notes

### Competing Interest Statement

The authors have declared no competing interest.

